# Identification of astrocytomas through serum protein fingerprint using MALDI-TOF MS and machine learning

**DOI:** 10.1101/2025.02.21.639567

**Authors:** Lucas C. Lazari, Janaina M. Silva, Priscila R. S. Donado, Sueli Mieko Oba Shinjo, Livia Rosa-Fernandes, Antonio Di Ieva, Suely K. N. Marie, Giuseppe Palmisano

## Abstract

Gliomas account for most brain malignancies, with astrocytomas being the most common subtype. Among these, glioblastoma (GBM) stands out as the most aggressive form, exhibiting a median survival time of just 15 months despite intensive therapy. Current diagnostic practices rely on magnetic resonance imaging (MRI) and histopathological analysis, which often necessitate invasive surgical sampling. This underscores the need for minimally invasive diagnostic tools capable of characterizing glioma progression and guiding treatment strategies. Advances in glioma classification have integrated histological and molecular markers, notably IDH1 mutations, which are prognostically significant, particularly in low-grade gliomas and in the previously defined “secondary GBM” (IDH-mutant astrocytoma grade 4). This study aimed to explore the potential of serum proteomics as a non-invasive diagnostic tool using MALDI-TOF mass spectrometry (MS) combined with machine learning techniques. We analyzed serum samples from 269 patients, employing machine learning models to differentiate between healthy individuals and astrocytoma patients. The MALDI-TOF MS approach achieved a balanced accuracy of 94.5% in distinguishing GBM patients from healthy controls. However, it showed limited efficacy in classifying tumor grades or determining IDH1 mutational status. Further investigation using bottom-up proteomics by GeLC-MS/MS identified potential biomarkers, such as transthyretin, previously associated with high-grade gliomas. These findings highlight the promise of MALDI-TOF MS in identifying serum-based biomarkers for astrocytoma diagnosis. While the results are promising, further validation in independent cohorts is essential to assess the clinical utility of these biomarkers for non-invasive glioma diagnostics and patient monitoring.

## Introduction

Gliomas account for approximately 80% of brain malignancies. Astrocytomas, which originate from astrocytic cells, are the most frequent type of glioma. Among these, glioblastoma (GBM, grade 4) is the most predominant subtype with the highest aggressiveness ^1,2^. With its median overall survival time of 15 months following combined treatment with radiotherapy and temozolomide, GBM is one of the most lethal human cancers ^3,4^. The most common diagnostic approach for GBM is radiographic characterization by using magnetic resonance imaging (MRI), with a definitive diagnosis based on histopathological analysis of the surgically-resected tumor ^5^. Risks related to surgical sampling make disease progression assessment suboptimal, therefore the development of minimally invasive diagnostic tools is paramount for a fast and accurate characterization of tumor’s evolution. According to the World Health Organization (WHO), gliomas are classified as grade 1, biologically benign tumor with low risk, generally surgically resectable; grades 2-3, low-grade glioma with differentiated cell pattern and better prognosis; and grade 4, high-grade glioma with undifferentiated cell pattern, highly aggressive and with poor prognosis.

A more recent glioma classification system was proposed based on in-depth tumor characterization, integrating histologic and molecular features ^6–8^. As a result of this integration of pheno- and genotypic features, gliomas are classified into two major categories: *IDH1*-wildtype (*IDH1*^wt^) and *IDH1*-mutant (*IDH1*^mut^) gliomas. *IDH* mutations occur most frequently at exon 4 codon 132 (R132H) and 80% of low-grade gliomas, grades 2 and 3, harbor this mutation ^9^. In GBMs, the *IDH* mutations are mainly found in secondary GBM (73%), whereas only 3.7% of primary GBM are *IDH*^mut 10^. The presence of *IDH* mutations correlates with favorable outcomes with a prolonged median survival. Low-grade *IDH1*^mut^ gliomas present 7-year overall survival, while primary *IDH1*^mut^ GBMs present 46 months of overall survival compared to 13 months in *IDH1*^wt^ GBMs ^11^. Therefore, the presence of *IDH* mutation impacts the overall survival and is considered a relevant prognostic marker for glioma diagnosis, prognosis and the establishment of treatment strategies ^10,12,13^.

Currently, the molecular diagnosis of gliomas can only be established by analyzing the tumor tissue resected during the surgical treatment. Cumulative efforts have been made to obtain biomarkers for classificatory diagnosis of gliomas with prognostic indications and therapeutic response, especially through a minimally invasive method, such as liquid biopsy or imaging techniques. Through the analysis of liquid biopsies, by drawing biofluids such as blood, saliva, urine, and/or cerebrospinal fluid (CSF), a plethora of molecules can be characterized as biomarkers. This approach has already been used for other tumor types, such as breast and lung cancer ^14–16^, and it has already been proved to possess both diagnostic and prognostic potential ^17–20^. In this context, for brain tumors, serum or plasma are the most used fluids in clinical testing due to their relative ease of acquisition through minimally invasive sampling, especially when compared to CSF. Indeed, a non-invasive early detection of gliomas in symptomatic patients could improve triage, thereby reducing the costs associated with biopsies and imaging. Additionally, these biomarkers could also be used to grade different gliomas for prognosis and treatment follow-ups. Among the analytical approaches developed to identify gliomas biomarkers, mass spectrometry-based techniques have shown some success ^21,22^.

The present study evaluated the performance of the MALDI-TOF MS combined with machine learning approach to analyze the serum proteomic fingerprint to discriminate between tumor and control patients, as well as low- and high-grade astrocytomas and their IDH mutational status in a cohort of over 197 astrocytoma and 72 control patients. To this end, we trained machine learning models on the MALDI-TOF MS data of sera collected from astrocytoma patients. This approach demonstrated to be capable of discriminating between healthy and GBM patients with a balanced accuracy of 94.5%, but failed to discriminate tumor grades or presence of IDH1 mutation. We further evaluated possible serum biomarkers that could be characteristic of GBM, thus we applied a large scale bottom-up proteomics approach focusing on serum proteins with low molecular weight (<15kDa) and detected the regulation of several proteins that might be associated with GBM, including proteins from APO family and Transthyretin, which has already been found in CSF and linked to high grade gliomas. This study shows the applicability of MALDI-TOF MS to select serum protein features followed by low molecular weight-directed protein identification to diagnose astrocytomas. The putative circulating and non-invasive biomarkers identified here need further validation in independent cohort to better appreciate their usefulness in clinical practice.

## Methods

### Cohort overview

The complete dataset consists of three groups: Control (72), AG2 (22), AG3 (20), and GBM (155). Two types of analysis were performed: Tumor (AG2 + AG3 + GBM) vs. Control comparison and pairwise comparison between the tumor classes (AG2 vs. AG3; AG3 vs. GBM; and AG2 vs. GBM).

Serum samples were collected during surgical procedures by the Neurosurgery Groups of Hospital das Clinicas, School of Medicine of University of Sao Paulo. Informed consent was obtained from each patient, and the study was approved by the national and local ethics committee under the number 64431622.3.0000.0068/5.899.401. The samples included frozen serum, collected upon surgical removal and immediately snap-frozen in liquid nitrogen. The mean age of AG2 patients was 37.2, with 45.5% females and 54.5% males; the mean age of AG3 patients was 35, with 40% females and 60% males; and the mean age of GBM patients was 53.5 years, with 36.1% females and 63.9% males. A total of 197 astrocytomas were analyzed and compared with 72 non-neoplastic control temporal lobe tissue collected from individuals submitted to epilepsy surgery with mean age of 56.2, being 65.3% females and 34.7% males. DNA was extracted from the frozen tissues by a standard method, and polymerase chain reaction (PCR) followed by DNA sequencing was applied to detect IDH1 mutation as described previously ^23^.

### Sample preparation

The sample preparation method was performed as described in Lazari et.al (2022) ^24^. A stage-tip micro-column was produced by inserting two C18 polymeric disks into p200 pipette tips. Column activation was conducted with 50 μl of 100% acetonitrile and conditioned with 100 μl of 0.1% TFA. Then, 1 µl of serum samples was diluted 10 times in an acid solution of TFA with a final concentration of 1%. Samples were carefully homogenized and added to the column. Then, 100 μl of 0.1% TFA was added to the column for a washing step and proteins were eluted with a solution of HCCA matrix (10 mg/mL in 50% acetonitrile, 25% water and 25% of 10% TFA) directly onto the MALDI plate. All steps except the elution were performed in a bench centrifuge at 2,000g for 2:30 min.

### Data acquisition

Samples were acquired in a MALDI-TOF Autoflex speed smartbeam mass spectrometer (Bruker Daltonics) using FlexControl software (version 3.3; Bruker Daltonics). Spectra were recorded in the positive linear mode (laser frequency, 500 Hz; extraction delay time, 390 ns; ion source 1 voltage, 19.5 kV; ion source 2 voltage, 18.4 kV; lens voltage, 8.5 kV; mass range, 2,400–20,000 D). Spectra were acquired using the automatic run mode to avoid subjective interference with the data acquisition. For each sample, 2,500 shots, in 500-shot steps, were summed. All spectra were calibrated by using Protein Calibration Standard I (Insulin [M+H]+ = 5,734.52, Cytochrome C [M+ 2H]2+ = 6,181.05, Myoglobin [M+ 2H]2+ = 8,476.66, Ubiquitin I [M+H]+ = 8,565.76, Cytochrome C [M+H]+ = 12,360.97, Myoglobin [M+H]+ = 16,952.31) (Bruker Daltonics).

### Data processing and machine learning analysis

The protocol used here was similar to the previous studies ^25–27^. Raw files were converted to the mzML format using the MSConvert function from the ProteoWizard suite ^28^, then processed using the MALDIQuant package ^29^. The spectra were square-root transformed, smoothed (method = SavitzkyGolay, halfWindowSize = 10), the base was corrected (method = TopHat) and normalized (method = Total Ion Current). Then, peaks were identified (halfWindowSize = 10 and SNR = 2), binned (method = strict and tolerance=0.003) and filtered keeping the ones present in at least 60% of the samples. Finally, data from both groups were merged and another binning step (method = strict and tolerance=0.003) was performed. The resultant peaks were used to plot the principal component analysis. For each pairwise analysis, a Shapiro-Wilk test for normality was performed in each peak, and the ones with a p-value < 0.05 were kept. Then, a Wilcoxon rank-sum test was performed to filter the most relevant peaks by keeping the ones with a p-value < 0.05. This step was performed to visualize if by selecting peaks it would be possible to find differences between the groups.

For the machine learning analysis, the normalized spectra were grouped into bins by summing all intensity values within 3 Da windows. We chose to use the entire binned spectra for the machine learning step because this approach simplifies preprocessing.

Selecting peaks separately for the training and testing sets would require their merge for peak alignment, which could introduce data leakage. Additionally, the availability of a substantial number of samples allowed us to work with an increased number of features. The training procedure was carried out by performing a 5-fold nested repeated 3-fold cross-validation (a final 75/25 train/test split). For each iteration of the outer loop, the dataset was split into training and test set, then oversampling was performed by using SMOTE, feature scaling was performed by fitting a StandardScaler in the training set, feature reduction was carried out by BorutaPy and then the training split was used to undergo a 3-fold cross-validation with random search for hyperparameter tunning. The best scoring model was tested using the test split. The mean balanced accuracy, specificity and sensitivity were calculated. A total of four models were tested (XGBoost, SVM, Random Forest and Multi Layer Perceptron), the highest performing model was reported.

The machine learning analysis was performed for the classification between health and tumor patients and for the classification of patients with and without IDH1-R132H.

### One dimensional SDS-PAGE

Proteins from all group’s samples were separated by one-dimensional gel electrophoresis using a 12% gel. The group distribution for this step was the following: 9 AG2 with IDH1 mutation, 9 AG2 without the mutation, 9 AG3 without mutation, 9 GBM with mixed mutation status and 9 controls. The gels were stained with Coomassie brilliant blue and scanned to identify differentially expressed protein bands. Bands corresponding to the molecular weight (MW) below 15,000 Da were excised for further analysis, this was done prioritizing the identification of low molecular weight proteins.

These bands were subjected to in-gel tryptic digestion, following the Shevchenko method, and analyzed using nanoflow LC-MS/MS. The nLC-MS/MS analysis was conducted on an Easy nano LC1000 HPLC system (Thermo Fisher Scientific) coupled with an LTQ Orbitrap Velos mass spectrometer (Thermo Fisher Scientific). Peptides were loaded onto a C18 EASY-column (2 cm × 5 µm × 100 µm, 120 Å pore; Thermo Fisher Scientific) at a flow rate of 300 nL/min with mobile phase A (0.1% formic acid), and separated using a C18 PicoFrit PepMap column (10 cm × 10 µm × 75 µm, 135 Å pore; New Objective) over 105 minutes with a linear gradient of 2–30% mobile phase B (100% acetonitrile with 0.1% formic acid). Peptides were ionized by electrospray, and the top 20 most intense precursor ions with a charge state of ≥2 were selected for fragmentation by collision-induced dissociation (CID) using a normalized collision energy of 35 and an activation time of 10 ms. The MS scan range was set to m/z 350–1,500, with an MS1 scan resolution of 60,000, an MS1 ion count target of 1 × 10^6^, and an MS2 ion count target of 3 × 10^4^. Raw data were submitted to ProteomeExchange (https://www.ebi.ac.uk/pride/).

### LC-MS/MS data analysis

nLC-MS/MS raw data were analyzed using MaxQuant for protein identification and label-free quantification. The raw files were searched against the Homo sapiens protein database, containing 20,397 reviewed protein sequences (UniProt, downloaded in October 2022). The database search was carried out using the trypsin as the proteolytic enzyme, allowing for two missed cleavages. Search parameters included a precursor ion tolerance of 10 ppm and a fragment ion mass tolerance of 0.5 Da. Carbamidomethylation of cysteine was specified as a fixed modification, while methionine oxidation was set as a dynamic modification. PSMs, peptides and proteins FDR was set to <1%.

All the identified proteins were filtered to remove contaminants and reverse-sorted, followed by a filter to select only those with at least one unique peptide. The LFQ intensities were then log2 transformed and filtered to have at least eight valid values in the total data set. Differentially regulated proteins were determined through a Kruskal-Wallis test and th.

## Results

The mass spectrum of all samples from Control and Tumor groups were used to generate a mean spectrum (**Figure 1A and B**), demonstrating a difference in peak intensity among the groups. The PCA analysis demonstrated that there is no clear separation between the tumor classes and IDH1 mutation status (**Figure 2**). The binning procedure to work with the whole spectra for model training was able to reduce the dimensionality of the data from approximately 32,000 to 4332 values, which were further reduced by BorutaPy during model training, all the selected features for the IDH1 mutation and tumor classifications for each fold are available in the **Supplementary Table 1 and 2**, respectively.

**Figure 1:**
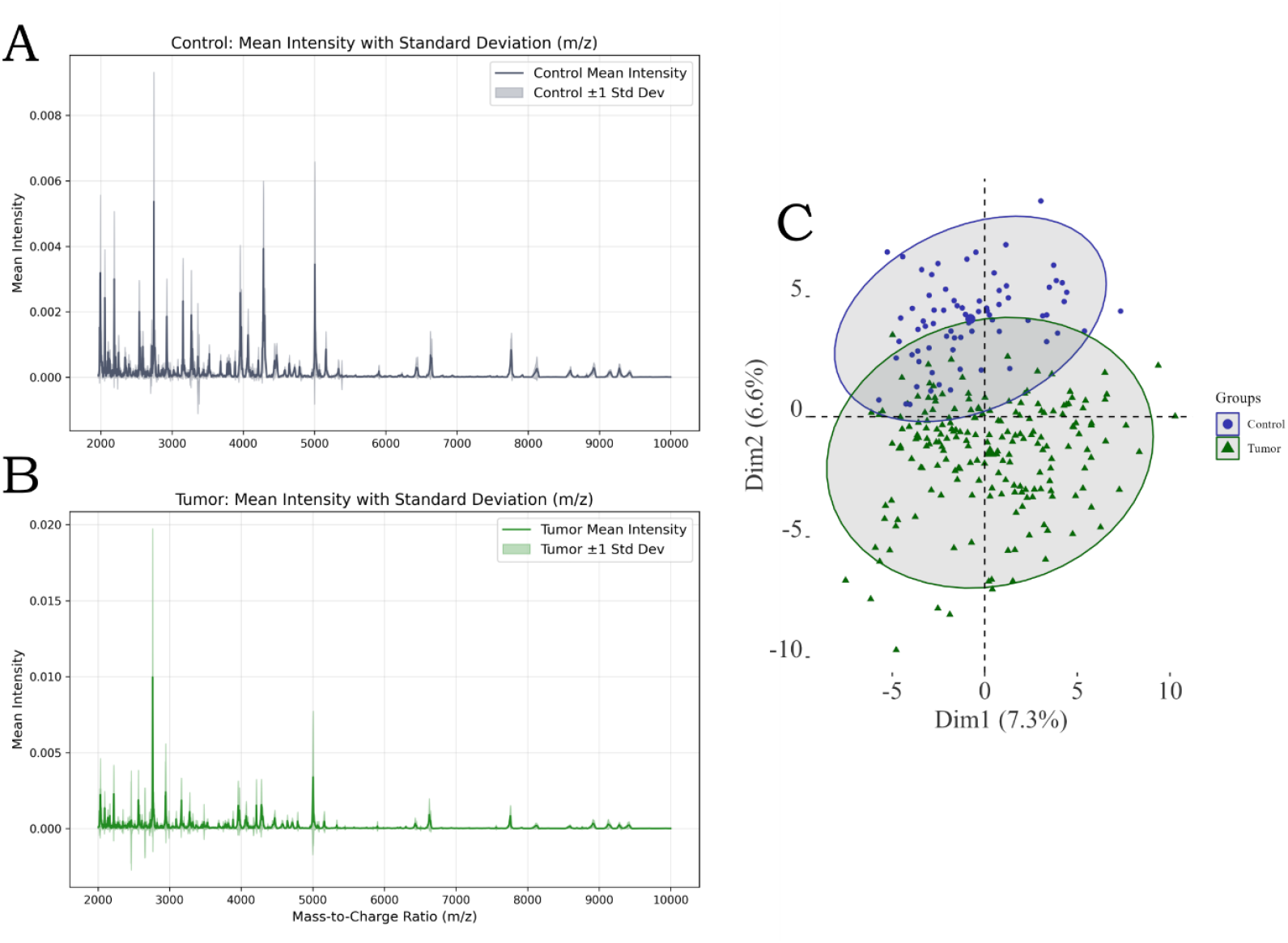
Mean spectra for the tumor (A) and control (B) groups with the standard deviation as a shaded area. PCA analysis (C) performed with the peaks identified in the spectra.

**Figure 2:**
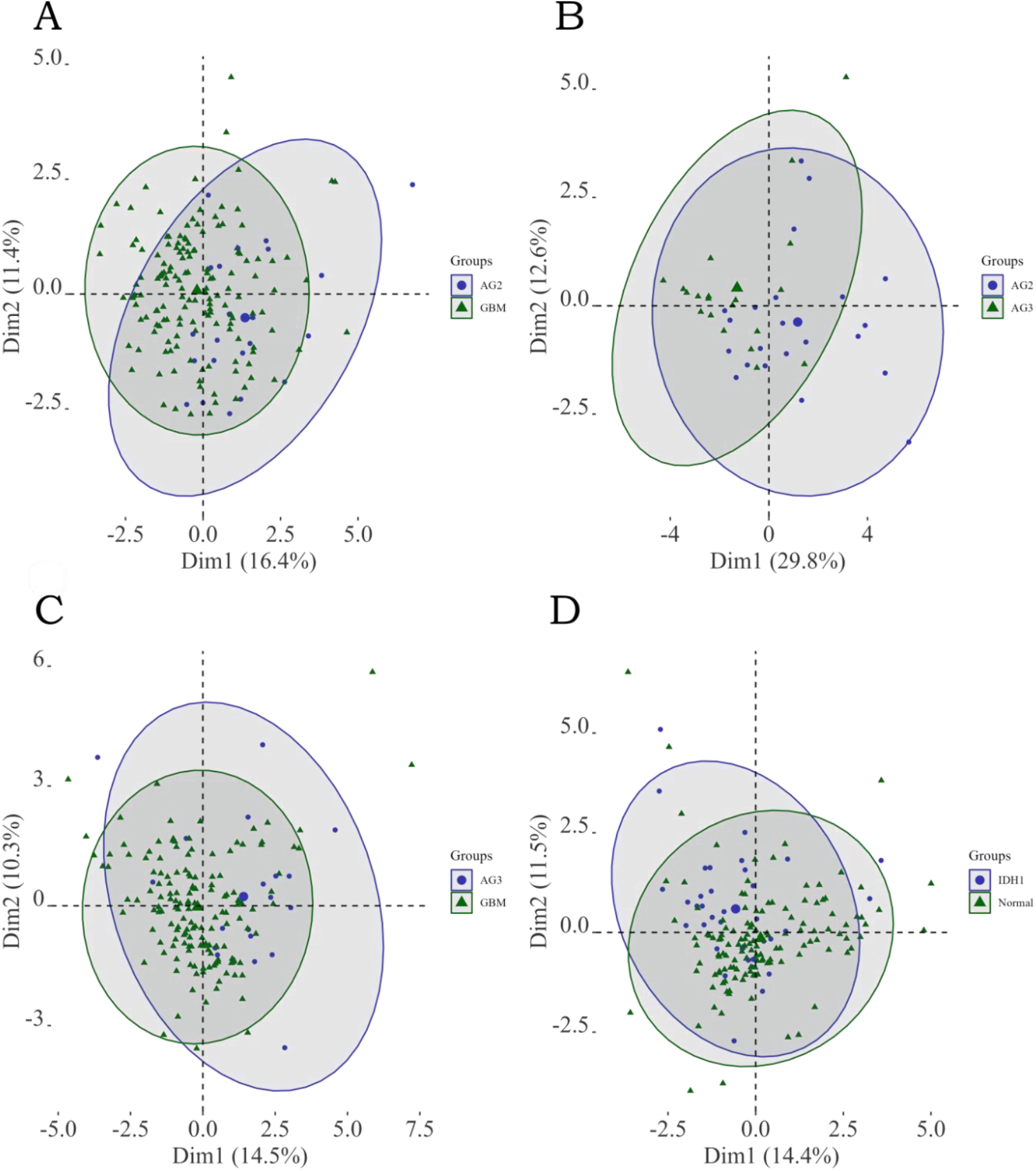
PCA analysis of the pair-wise comparisons between the tumor classes (A, B and C) and IDH1 mutation status (D).

**Figure 3:**
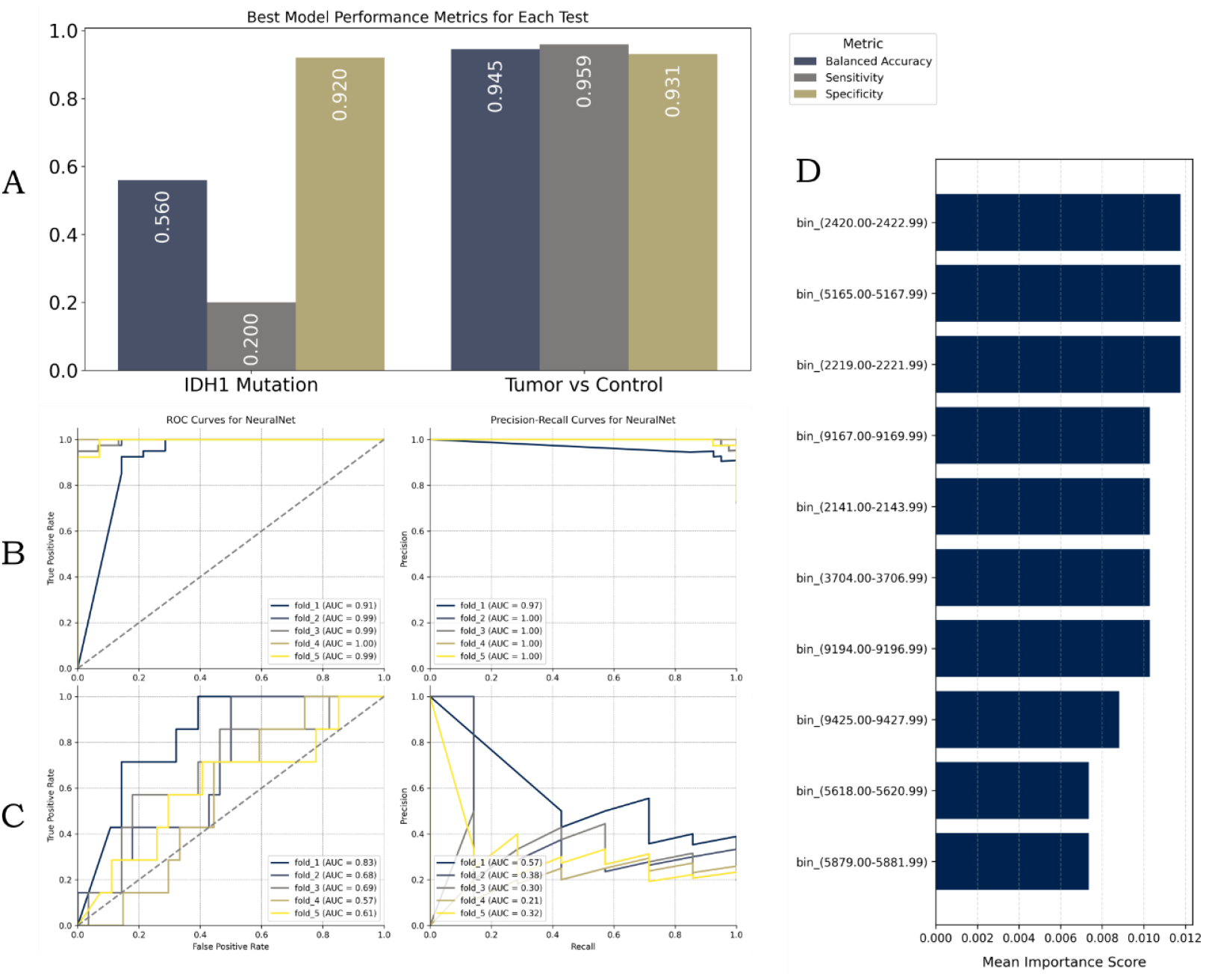
Multi-Layer Perceptron (NeuralNet) metrics for both IDH1 mutation status and tumor classification (A). Receiver Operant Curves and Precision Recall Curves for tumor classification (B). Receiver Operant Curves and Precision Recall Curves for IDH1 mutation status (C). Top 10 most important features for tumor classification accordingly to permutation importance (D).

All the training models for the tumor versus controls classification achieved over 87% balanced accuracy, with the highest scoring model (Multi Layer Perceptron) scoring the highest with 94.5% balanced accuracy (**Figure 2A**). For the IDH1 mutation status classification, all the models failed to discriminate the groups, being the highest balanced accuracy 56% (**Figure 2A**). The ROC and PR curves for the best model are reported in **Figure 2B**. All metrics are available at the **Supplementary Table 3** for the IDH1 mutation and tumor classification, **Supplementary Tables 4 and 5** contains all hyperparameters for each fold for the IDH1 mutation and tumor classification respectively. For the tumor versus control comparison we searched for the most important features for classification by using permutation importance from sklearn.inspection in the highest scoring model. The top 10 most important features are reported in Figure 2C.

To identify proteins differentially regulated in astrocytomas, we applied a LC-MS/MS approach on the serum proteins separated on a 1D SDS-PAGE and being below 15kDa, since most of the important features accordingly to permutation entropy were within this range. A principal component analysis (PCA) was also performed using all proteins identified (**Figure 4A**), but no clear separation of the groups was observed. However, the data showed differentially regulated proteins that could be possible markers to differentiate between groups (**Figure 4B**). More interestingly, the heatmap showed that TTR, GC (Vitamin D Binding Protein) and PRG4 (Proteoglycan 4) proteins are more abundant in GBM when compared to the other tumor grades.

**Figure 4:**
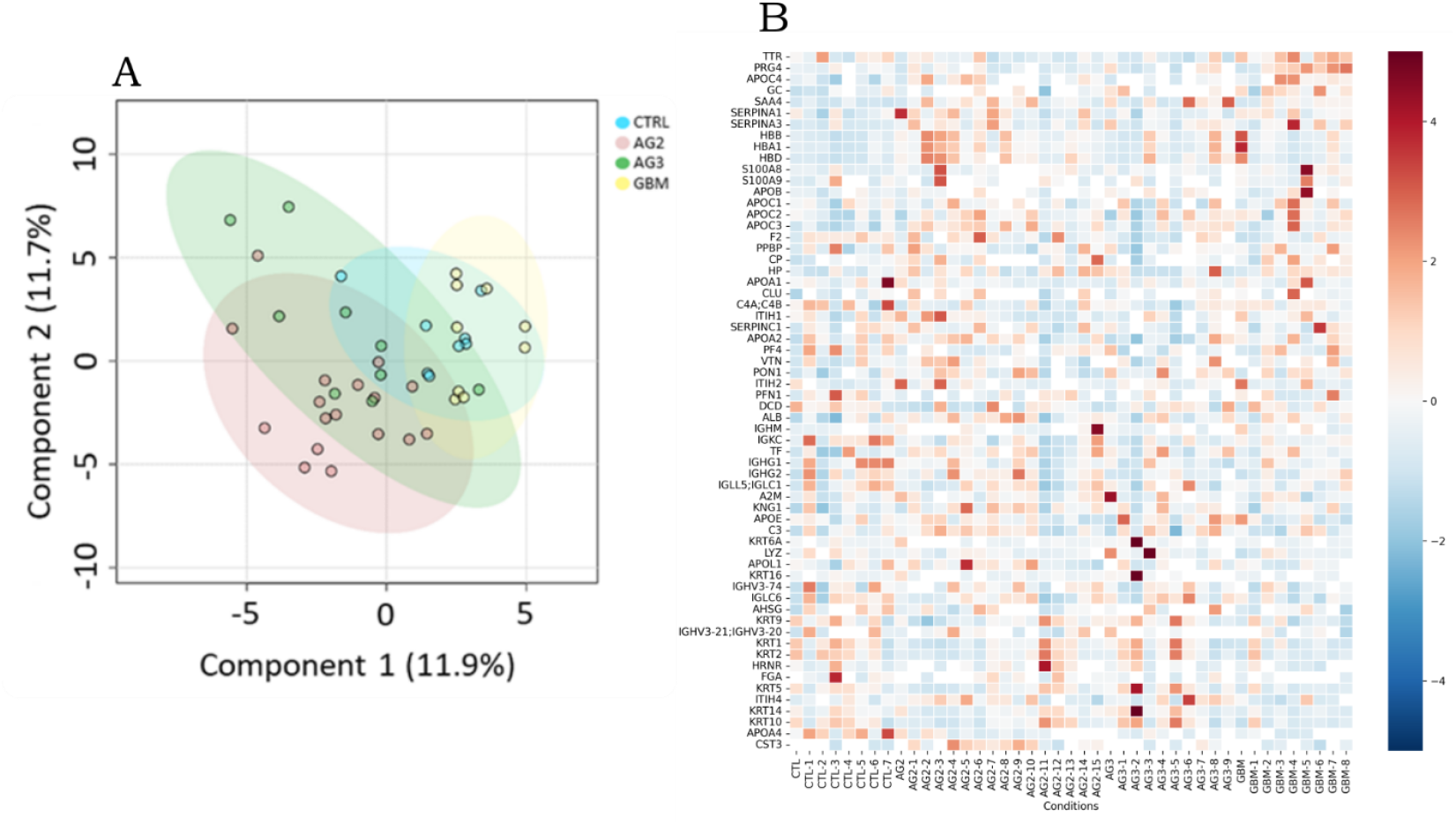
(A) Principal Component Analysis of all identified proteins. (B) Heatmap of the differentially regulated proteins from the LC-MS/MS analysis of low molecular weight proteins in GBM.

## Discussion

Despite the recent advances in medicine the survival rate of GBM patients still ranges from 14 to 17 months ^30^. The therapeutic challenges highlight the need for the development of early and precise diagnosis methods, as early intervention could potentially lead to tumor progression mitigation enhancing patient survivability. We demonstrated that the combination of MALDI-TOF MS protein profiling of serum samples can discriminate non-tumor and tumor patients. The use of proteomics to search for GBM biomarkers is already well-studied in the literature. Many protein biomarkers were already reported for diagnosis and prognosis purposes, and different mass spectrometry approaches were tested in GBM samples. Among the findings, plasma proteins discovered using SWATH-MS, such as CRP, C9, and LRG1 were reported to have a positive correlation with tumor size, while GSN protein was correlated with survival time ^31^. A spatial resolved microproteomics of tumor samples using MALDI-MSI, SpiderMass and NanoLC-MS/MS analysis was also able to identify a set of biomarkers that are indicative of unfavorable prognosis (ALCAM, ANXA11, and AltProt IP_652563 ^32^. The serum proteins S100A8, S100A9, and CXCL4 identified using a combination of SELDI-TOF MS and LC-MS/MS were reported to serve as potential biomarkers capable of discriminating GBM patients from controls ^33^. In another study, the MALDI-TOF MS profile of human cancer cells revealed specific peaks between the spectra of U118, U138, U87, and U373 cell lines with different resistance to treatment ^34^.

The MALDI-TOF mass spectrometry method employed in this work was initially used to select serum protein features that discriminated the different pathological conditions. Although it is limited in this aspect, the methods for sample processing and data acquisition are simplified, being faster and low-cost when compared with other mass spectrometry techniques. Machine learning then makes it possible to develop a diagnosis method based in the MALDI-TOF protein profiling signature, allowing patient classification without the need for protein identification.

MALDI-TOF MS has been used in brain tumors in imaging and coupled with 2D-electrophoresis for spot identification mainly in tumor tissues ^32,35–39^. Another study applied MALDI-TOF MS to discriminate U87-MG GBM cell before and after treatment with sodium phenylbutyrate and all-trans retinoic acid ^40^. Changes in lipid profile discriminated the different cells. Besides its clinical application in clinical microbiology, MALDI-TOF MS can be used to profile circulating markers in cancer, MALDI-TOF MS detection of circulating biomarkers has been implemented in a commercially available test for lung cancer, VeriStrat, originally developed to identify non-small cell lung cancer (NSCLC) patients’ non-responders to second line gefitinib treatment ^41^. This assay is based on eight MALDI-TOF MS peaks, partially associated to proteoforms of serum amyloid A protein, obtained from serum or plasma that can serve as predictive markers to classify NSCLC patients as good or poor responders to EGFR TKI therapy ^42^. Few studies have applied SELDI-TOF and MALDI-TOF MS to profile the circulating proteome in gliomas. Comparing serum samples from astrocytoma patients with samples from control subjects, a panel of 7 mass spectrometric peaks were significantly regulated and showed high accuracy in discriminating these conditions using protein chip array and SELDI-TOF acquisition ^43^. Another study used a similar experimental approach combined with artificial neural network (ANN) algorithm to identify 22 serum peptides for the identification of glioma versus brain benign tumors with high accuracy ^44^. Another approach used magnetic reversed-phase derivatized particles combined with MALDI-TOF MS to profile, in an automated way, the serum of glioma patients and healthy controls. A pattern of 274 peptide masses achieved high accuracy in discriminating these conditions ^45^. One study analyzed the serum proteome profile of GBM patients using MALDI-TOF MS by analyzing the serum proteins/peptides absorbed on cation exchange magnetic beads. The comparison between GBM patients and healthy subjected revealed a combination of 11 MALDI-TOF MS peaks that discriminated the two groups with high accuracy. However, the same was not observed while separating between grades ^46^. Another study used SELDI-TOF MS to compare the serum proteome profile in GBM patients and healthy subjects. Furthermore, the identification of proteins with a MW < 28kDa, similar to the range detected in the SELDI-TOF, was performed using 1D gel electrophoresis coupled with LC-MS/MS. Three proteins S100A8, S100A9 and CXCL4 were validated as potential biomarkers using ELISA and S100A9 and CXCL4 confirmed by western blotting to be highly expressed in tumoral compared to peritumoral GBM tissues ^33^. Due to that, the study presented here shows the application of MALDI-TOF MS and computational approaches to diagnose astrocytomas.

Complementary to tumor identification, the assessment of tumor progression is also desirable to ensure the proper treatment methods. The identification of IDH1-mutant gliomas is an important indicative for GBM prognosis ^12^. Through a PCA analysis we demonstrated that the protein profile of AG2 tumors patients differs from AG3 and GBM patients. This indicates that the MALDI-TOF protein profiling may be able to separate low-grade and high-grade tumors. The results also demonstrate that AG3 and GBM have a very similar MALDI-TOF MS profile. It is important to note that the number of samples of the AG2 and AG3 groups were limited compared to the GBM group, therefore, to consolidate this finding a higher number of samples from these groups would be necessary.

Although a difference between low-grade and high-grade tumors was observed, it was not possible to differentiate IDH1 mutations from non-mutated individuals using the methods and the biofluid (serum) employed in this work. The alterations caused by IDH1 mutation includes production of high levels of 2-hydroxyglutaric acid (2-HG), up-regulation of vascular endothelial growth factor (VEGF) and production of high levels of hypoxia-inducible factor-1α (HIF-1α) ^13^. It is possible that the methods for protein purification performed in this work were not sensitive enough to capture alterations in one of these molecules, especially 2-HG, which is a metabolite, therefore elusive in our analysis. The identification of 2-HG in tumor sections by MALDI-TOF and MALDI-MSI was already reported in the literature ^47–49^; however, the identification of this molecule in serum samples needs to be optimized. Therefore, further optimization and standardization of MALDI-TOF MS protocols are needed to ensure the robustness and reliability of serum and biomarker discovery.

A recent study evaluated the circulating proteome in serum using the proximity-ligation assay in a small cohort of patients with GBM, anaplastic astrocytoma, anaplastic oligodendrogliomas and meningioma was used as a control ^50^. GFAP and FABP4 were identified as discriminant biomarkers for GBM and meningiomas. Another study evaluated the autoantibody profile of gliomas patients using an array of 17000 full-length human proteins. The protein recognized by serum autoantibodies were able to distinguish between gliomas and healthy subjects with good sensitivity and specificity. However, there was not a clear separation between grades and IDH1 mutational status, showing how hard it is too discriminate these conditions ^51^.

To identify the low molecular weight protein markers responsible for discriminating the gliomas, we performed a GeLC-MS approach. We also found that Transthyretin (TTR) appears to be more abundant in GBM when compared to the other tumor grades. This protein has already been detected up-regulated in CSF ^52^ and was linked to high grade gliomas ^53^. Interestingly we detected this protein regulation using serum samples, highlighting its capabilities as a possible marker for grade differentiation. The LC-MS/MS data also indicates that GC protein might be a characteristic of GBM patients, high abundance of GC proteins may reduce the bioavailability of circulating vitamin D by tightly binding to it, limiting its “free” active form accessible to tissues. A study found cancer survival advantages only in individuals with the Gc2 isoform, where lower GC protein levels allow more unbound vitamin D to exert protective effects. Thus, individuals with Gc1 isoforms may require higher systemic vitamin D concentrations to achieve comparable bioactive levels ^54^.

## Conclusions

Despite recent advances in medicine, the survival rate for GBM patients remains low, emphasizing the urgency for refined diagnostic techniques and early intervention. This research underscores the potential of MALDI-TOF MS protein profiling in serum samples as a promising tool for distinguishing between healthy and individuals with tumor. The cost-effectiveness, speed, and simplified protocols alongside the application of machine learning turns MALDI-TOF MS a good alternative to other mass spectrometry methods for the development of diagnostic and prognostic tools for gliomas. Furthermore, while the study indicates a distinction between low-grade and high-grade tumors, it could not differentiate between IDH1 mutations. The inability to detect alterations caused by the IDH1 mutation, particularly in molecules like 2-HG, suggests that other sample preparation methods will be needed to capture those alterations. Future efforts should focus on enhancing these methodologies and expanding the sample size for more accurate and conclusive results validating the potential biomarkers identified here.

## Funding

We are grateful for the financial support provided by the São Paulo Research Foundation (FAPESP, grants processes n° 2018/18257-1 (GP), 2018/15549-1 (GP), 2020/04923-0 (GP), 2021/00140-3 (JMDS), 2020/02988-7 (SKNM, SMOS); by the Conselho Nacional de Desenvolvimento Científico e Tecnológico (“Bolsa de Produtividade” (SMOS, SKNM and GP); by Fundação faculdade de Medicina (FFM-SKNM); and by Fundação Coordenação de Aperfeiçoamento de Pessoal de Nível Superior (CAPES, processes 88887.510020/2020-00 and 88887.884790/2023-00).

## Notes

### Competing Interest Statement

The authors have declared no competing interest.

### Summary of Updates

There are no changes in the content of the article. The main and unique change was in the authors order, since there was a mistake during the submission process.

